# Cryopreservation of Ghagus chicken semen: effect of cryoprotectants diluents and thawing temperature

**DOI:** 10.1101/2020.05.16.099333

**Authors:** Shanmugam Murugesan, Ramkrishna Mahapatra

**Affiliations:** ICAR-Directorate of Poultry Research, Rajendranagar, Hyderabad-30, Telangana, India

**Keywords:** Chicken, Cryoprotectant, Diluent, Fertility, Semen cryopreservation

## Abstract

The present study evaluated the effects of cryoprotectants, semen diluents and thawing temperature during Ghagus chicken semen cryopreservation. Four different experiments were conducted; Experiment 1- semen was cryopreserved using 6% dimethylacetamide (DMA) and 2% dimethylsulfoxide (DMSO) in Sasaki diluent (SD) and Lake and Ravie diluent (LRD), Experiment 2 and 3- semen was cryopreserved using 8% Ethylene Glycol (EG) in SD, LRD and Red Fowl Extender (RFE), Experiment 4- semen was cryopreserved using 6% dimethylformamide (DMF) in SD, LRD and Beltsville Poultry Semen Extender (BPSE). Semen was cryopreserved in 0.5 ml French straws. Thawing was done at 5°C for 100 sec in ice water in Experiments 1, 2 and 4, whereas in Experiment 3 thawing was done at 37°C for 30 sec. The post-thaw sperm motility, live sperm and percent acrosome intact sperm were significantly (*P*<0.05) lower in cryopreserved samples in all the experiments. No fertile eggs were obtained from cryopreserved samples in Experiments 1 and 2, except for 8% EG RFE treatment where the fertility was 0.83%. In Experiments 3 and 4, highest fertility was obtained in LRD treatment 48.12 and 30.89% respectively. In conclusion, using cryoprotectant EG (8%) and thawing at 37°C for 30 sec, and DMF (6%) resulted in acceptable level of fertility in Ghagus chicken. Though the diluents influenced post-thaw in vitro semen parameters the fertility was not affected. In addition, results indicated that thawing temperature may be a critical stage in the cryopreservation protocol.

## 1 INTRODUCTION

Semen cryopreservation is an approach for long term *ex-situ* conservation of genetic resources because of the cost effectiveness and simplicity of the technique (Silversides et al., 2012). The fertilizing ability and viability of cryopreserved sperm is different between chicken breeds requiring standardization of breed or line specific semen cryopreservation protocol (Blesbois 2011; Long 2006). Ghagus is an important native chicken breed of India with native tract in Karnataka state. These birds are of medium size with good mothering ability and broodiness character (Haunshi et al., 2015). Efforts are being made for *ex-situ* conservation of this breed of chicken.

The diluent used during semen cryopreservation should essentially help in maintaining the structure and functional capacity of the sperm during and after cryopreservation. Few studies have compared the effect of diluents/extenders during chicken semen cryopreservation (Nabi et al., 2016; Rakha et al., 2016; Roushdy et al., 2014; Shanmugam and Mahapatra, 2019). In addition to energy and buffering components in diluents used for semen cryopreservation substances such as polyvinylpyrrolidone and glutamic acid are included that supports the sperm during cryopreservation procedure. A diluent with minimal components that may support sperm during liquid storage may not be of use during cryopreservation (Shanmugam and Mahapatra, 2019).

The cryoprotectant used for semen cryopreservation by itself profoundly influences the post-thaw semen parameters and fertility (Long, 2006). Glycerol is the least toxic and effective poultry semen cryoprotectant, however, concentration above 1% during insemination produces contraceptive effect (Lake et al., 1980) necessitating it to be removed before insemination. Various other cryoprotectants such as dimethylsulfoxide (DMSO), ethylene glycol and those belonging to amide group such as dimethylacetamide, dimethylformamide and methylacetamide were used for cryopreserving poultry semen and the semen can be used for insemination without removal of the cryoprotectant.

Semen cryopreservation protocol for preserving Ghagus breed is not available. This experiment aimed to develop a semen cryopreservation protocol for Ghagus chicken and study the effect of different cryoprotectants, semen diluents and thawing temperature.

## 2 MATERIALS AND METHODS

### 2.1 Experimental birds and husbandry

The experiment was conducted at the experimental poultry farm of ICAR-Directorate of Poultry Research, Hyderabad, India. The birds were housed in individual cages in an open-sided house and feed and water was available *ad libitum* throughout the experimental period. The study consisted of four independent experiments and was carried out following the approval of the Institutional Animal Ethics Committee.

### 2.2 Experimental procedures

#### Experiment 1

Semen from fifteen Ghagus rooster (32 weeks age) was collected by abdominal massage (Burrows and Quinn, 1937), pooled and used for the experiment. An aliquot of pooled semen was evaluated for sperm concentration, progressive motility, live and abnormal sperm. Semen was cryopreserved using 6% DMA and 2% DMSO in two diluents, Sasaki diluent (D (+)-glucose- 0.2 g, D (+)-trehalose dehydrate- 3.8 g, L-glutamic acid, monosodium salt- 1.2 g, Potassium acetate- 0.3 g, Magnesium acetate tetrahydrate- 0.08 g, Potassium citrate monohydrate- 0.05 g, BES- 0.4 g, Bis-Tris- 0.4 g in 100 ml distilled water, final pH 6.8; Sasaki et al., 2010) and Lake and Ravie (LR) diluent (sodium glutamate 1.92 g, glucose 0.8 g, magnesium acetate 4H_2_O 0.08 g, potassium acetate 0.5 g, polyvinylpyrrolidone [relative molecular mass (Mr) = 10 000] 0.3 g and double distilled water 100 ml, final pH 7.08, osmolality 343 mOsm/kg; Lake and Ravie, 1984). The semen mixed with the cryoprotectant with final sperm concentration 2000 × 10^6^/ml was immediately loaded into 0.5 ml French straws and sealed with polyvinyl alcohol powder. The straws were placed 4.5 cm above liquid nitrogen (LN_2_) on a styrofoam raft floating on LN_2_ in a thermocol box and exposed to nitrogen vapours for 30 minutes. The straws were then plunged into LN_2_ and stored till further use. Semen straws were stored for a minimum of seven days before evaluation. Post-thaw evaluation and insemination of semen was done after thawing at 5°C for 100 sec in ice water (Sasaki et al., 2010). The samples were evaluated on seven different occasions for progressive sperm motility, live sperm, abnormal sperm, and intact sperm acrosome. Thawed semen was inseminated per vagina into 33 weeks old Ghagus hens (13 hens/treatment) using a dose of 200 million sperm/0.1 ml volume. Insemination was repeated three times at three days interval. Freshly collected and inseminated semen served as control. Eggs were collected from the second day onwards after first insemination and stored in cold chamber (15°C) till incubation. Eggs were candled on the 18^th^ day of incubation for embryonic development. Infertile eggs were broke open for examination and confirmation of the absence of embryonic growth. Hatchability results were obtained on 21^st^ day of incubation.

#### Experiment 2

Ghagus (41 weeks age) semen was cryopreserved similar to Experiment 1 with few modifications. Ethylene glycol at 8% final concentration was used as cryoprotectant and three semen diluents namely Sasaki diluent, Lake and Ravie diluent and Red Fowl Extender (RFE; Fructose-1.15 g, Sodium glutamate-2.1 g, Polyvinylpyrrolidone- 0.6 g, Glycine- 0.2 g, Potassium acetate- 0.5 in 100 ml distilled water, final pH 7, osmolality 380 mOsm/kg; Rakha et al., 2016) were evaluated. Post-thaw semen was evaluated on six occasions for progressive sperm motility, live sperm, abnormal sperm, and intact sperm acrosome. Thawed semen was inseminated into 46 weeks old Ghagus hens (15 hens/treatment) using a dose of 200 million sperm in 0.1 ml volume. Insemination was repeated three times at four days interval. Insemination using fresh semen served as control. Post insemination eggs were incubated and data on fertility parameters obtained as in Experiment 1.

#### Experiment 3

In this experiment Ghagus (54 weeks age) semen was cryopreserved and evaluated similar to Experiment 2 except that the semen straws were thawed at 37°C for 30 sec. Thawed semen was inseminated into 54 weeks old Ghagus hens (15 hens/treatment) using a dose of 200 million sperm in 0.1 ml volume. Insemination was repeated three times at four days interval. Post insemination eggs were incubated and data on fertility parameters obtained as in Experiment 1.

#### Experiment 4

Ghagus (54 weeks age) semen was cryopreserved similar to Experiment 1 with few modifications. Dimethylformamide at 6% final concentration was used as cryoprotectant and three semen diluents namely Sasaki diluent, Lake and Ravie diluent and Beltsville Poultry Semen Extender (BPSE; Fructose-0.3 g, Potassium citrate-0.0384 g, Sodium glutamate-0.5202 g, Magnesium chloride- 0.0204 g, di-Potassium hydrogen phosphate- 0.762 g, TES- 0.317 g, Potassium di-hydrogen phosphate- 0.039 g, Sodium acetate- 0.258 g in 100 ml distilled water, final pH 7.3, osmolality 330 mOsm/kg; Rakha et al., 2016) were evaluated. Post-thaw semen was evaluated on six occasions for progressive sperm motility, live sperm, abnormal sperm, and intact sperm acrosome. Thawed semen was inseminated into 57 weeks old Ghagus hens (20 hens/treatment) using a dose of 200 million sperm in 0.1 ml volume. Insemination was repeated three times at four days interval. Insemination using fresh semen served as control. Post insemination eggs were incubated and data on fertility parameters obtained as in Experiment 1.

### 2.3 Semen quality assays

#### Sperm motility

Sperm motility was subjectively scored as percentage of progressively motile sperm after placing a drop of diluted semen on a Makler chamber and examined under 20x magnification.

#### Live and abnormal sperm

The live and abnormal sperm percent was estimated by differential staining using Eosin-Nigrosin stain (Campbell et al. 1953). Semen smear prepared by mixing one drop of semen with two drops of Eosin-Nigrosin stain was air dried and observed under high power (1000x). All pink stained and partially stained sperm were considered dead and unstained sperm as live. The percentage of live sperm was determined by counting at least 200 sperm. The same slides were used for estimating the percentage of abnormal sperm that was showing different morphological abnormalities.

#### Intact sperm acrosome

The intact acrosome in sperm was assessed as described by Pope et al. (1991). Briefly, 10 μl of diluted semen was mixed with 10 μl of stain solution (1% (wt/vol) rose Bengal, 1% (wt/vol) fast green FCF and 40% ethanol in citric acid (0.1 M) disodium phosphate (0.2 M) buffer (McIlvaine’s, pH 7.2-7.3) and kept for 70 sec. A smear from the mixture was made on glass slide, dried and examined under high magnification (1000x). The acrosomal caps were stained blue in acrosome-intact sperm and no staining in the acrosome region of acrosome reacted sperm. A minimum of 200 sperm were counted in each smear sample for calculating the percent acrosome intact sperm.

### 2.4 Statistical analysis

Data were analyzed using SAS 9.2 software and *P*<0.05 was considered significant. Statistical analyses of semen parameters were performed by one-way ANOVA with Tukey’s post hoc test. Data with percent values were arcsine transformed before analysis.

## 3 RESULTS

The post-thaw sperm motility and live sperm percent were significantly (*P*<0.05) lower in cryopreserved samples in all the experiments (Tables 1–4). Cryopreservation did not affect the percent abnormal sperm in first three experiments and in Experiment 4 samples cryopreserved in LR and BPSE diluents had significantly (*P*<0.05) higher abnormal sperm. Intact sperm acrosome was significantly (*P*<0.05) lower in cryopreserved sperm except for the 8% EG SD treatment in Experiment 3. No fertile eggs were obtained from cryopreserved samples in Experiment 1 and 2, except for 8% EG RFE treatment where the fertility was 0.83% (Table 2). The percent fertility after insemination with cryopreserved semen was significantly (*P*<0.05) lower in all the treatments of Experiments 3 and 4 except for 8% EG LR treatment where it was similar to that of control (Tables 3 and 4). No difference in percent hatchability was observed between the treatments in Experiments 3 and 4.

**TABLE 1.** Effect of cryoprotectants and semen diluents on post thaw semen and fertility parameters in cryopreserved Ghagus semen.

| Parameters | Control | 6% DMA SD | 2% DMSO SD | 6% DMA LR | 2% DMSO LR |
| --- | --- | --- | --- | --- | --- |
| Progressive sperm motility (%) | 64.29 ± 1.7<sup>a</sup> | 21.43 ± 1.43<sup>b</sup> | 12.14 ± 1.01<sup>c</sup> | 17.86 ± 1.01<sup>b</sup> | 9.28 ± 0.71<sup>c</sup> |
| Live sperm (%) | 71.83 ± 2.92<sup>a</sup> | 34.93 ± 1.69<sup>b</sup> | 22.66 ± 0.95<sup>c</sup> | 34.46 ± 2.13<sup>b</sup> | 21.23 ± 1.51<sup>c</sup> |
| Abnormal sperm (%) | 2.07 ± 0.63 | 2.13 ± 0.62 | 1.51 ± 0.22 | 2.51 ± 0.35 | 1.04 ± 0.18 |
| Acrosome intact sperm (%) | 99.57 ± 0.2<sup>a</sup> | 4.57 ± 1.21<sup>c</sup> | 94.85 ± 0.55<sup>b</sup> | 4.14 ± 0.74<sup>c</sup> | 91.61 ± 0.76<sup>b</sup> |
| Fertility (%) | 85.76 ± 5.47<sup>a</sup> | 0<sup>b</sup> | 0<sup>b</sup> | 0<sup>b</sup> | 0<sup>b</sup> |
| Hatchability on FES (%) | 98.18 ± 1.81<sup>a</sup> | 0<sup>b</sup> | 0<sup>b</sup> | 0<sup>b</sup> | 0<sup>b</sup> |
| No. of eggs incubated | 43 | 34 | 35 | 29 | 52 |

Values given are Mean±SE.
Figures bearing different superscripts in a row differ significantly ( $P<0.05$ ).
FES- Fertile egg set

**TABLE 2.** Effect of ethylene glycol and semen diluents on post-thaw semen and fertility parameters in Ghagus chicken thawed at 5°C.

| Parameters | Control | 8% EG SD | 8% EG LR | 8% EG RFE |
| --- | --- | --- | --- | --- |
| Progressive sperm motility (%) | 62.5 ± 1.12<sup>a</sup> | 20.0 ± 1.3<sup>b</sup> | 18.33 ± 1.05<sup>bc</sup> | 15.0 ± 1.29<sup>c</sup> |
| Live sperm (%) | 74.22 ± 1.64<sup>a</sup> | 23.72 ± 1.12<sup>b</sup> | 26.52 ± 1.54<sup>b</sup> | 22.72 ± 1.43<sup>b</sup> |
| Abnormal sperm (%) | 2.7 ± 0.23 | 2.15 ± 0.19 | 3.02 ± 0.40 | 2.58 ± 0.42 |
| Acrosome intact sperm (%) | 99 ± 0.37<sup>a</sup> | 94.0 ± 0.86<sup>b</sup> | 92.33 ± 0.92<sup>b</sup> | 92.83 ± 1.08<sup>b</sup> |
| Fertility (%) | 71.63 ± 4.81<sup>a</sup> | 0<sup>b</sup> | 0<sup>b</sup> | 0.83 ± 0.83<sup>b</sup> |
| Hatchability on FES (%) | 98.81 ± 1.19<sup>a</sup> | 0<sup>b</sup> | 0<sup>b</sup> | 0<sup>b</sup> |
| No. of eggs incubated | 82 | 85 | 108 | 86 |

Values given are Mean±SE.
Figures bearing different superscripts in a row differ significantly ( $P<0.05$ ).
FES- Fertile egg set

**TABLE 3.** Effect of ethylene glycol and semen diluents on post-thaw semen and fertility parameters in Ghagus chicken thawed at 37°C.

| Parameters | Control | 8% EG SD | 8% EG LR | 8% EG RFE |
| --- | --- | --- | --- | --- |
| Progressive sperm motility (%) | 64.16 ± 1.54<sup>a</sup> | 17.5 ± 1.12<sup>b</sup> | 17.5 ± 1.12<sup>b</sup> | 11.67 ± 1.05<sup>c</sup> |
| Live sperm (%) | 80.17 ± 3.30<sup>a</sup> | 26.0 ± 2.03<sup>b</sup> | 24.67 ± 2.0<sup>b</sup> | 20.33 ± 0.71<sup>b</sup> |
| Abnormal sperm (%) | 1.48 ± 0.27 | 1.87 ± 0.24 | 1.93 ± 0.18 | 1.8 ± 0.17 |
| Acrosome intact sperm (%) | 97.17 ± 0.31<sup>a</sup> | 84.58 ± 3.84<sup>ab</sup> | 69.98 ± 8.11<sup>b</sup> | 62.73 ± 9.60<sup>b</sup> |
| Fertility (%) | 81.23 ± 5.39<sup>a</sup> | 18.39 ± 8.89<sup>b</sup> | 48.12 ± 11.1<sup>ab</sup> | 38.29 ± 10.27<sup>b</sup> |
| Hatchability on FES (%) | 72.53 ± 4.18 | 69.12 ± 14.98 | 68.98 ± 6.4 | 58.1 ± 9.85 |
| No. of eggs incubated | 82 | 101 | 84 | 111 |

Values given are Mean±SE.
Figures bearing different superscripts in a row differ significantly ( $P<0.05$ ).
FES- Fertile egg set

**TABLE 4.** Effect of dimethylformamide and semen diluents on post-thaw semen and fertility parameters in Ghagus chicken.

| Parameters | Control | 6% DMF SD | 6% DMF LR | 6% DMF BPSE |
| --- | --- | --- | --- | --- |
| Progressive sperm motility (%) | 66.43 ± 1.43<sup>a</sup> | 18.57 ± 1.8<sup>b</sup> | 20.0 ± 2.44<sup>b</sup> | 15.0 ± 1.54<sup>b</sup> |
| Live sperm (%) | 79.47 ± 4.48<sup>a</sup> | 26.0 ± 1.73<sup>b</sup> | 23.81 ± 2.79<sup>b</sup> | 23.53 ± 1.06<sup>b</sup> |
| Abnormal sperm (%) | 1.32 ± 0.15<sup>b</sup> | 2.32 ± 0.34<sup>ab</sup> | 3.44 ± 0.54<sup>a</sup> | 3.34 ± 0.5<sup>a</sup> |
| Acrosome intact sperm (%) | 97.77 ± 0.14<sup>a</sup> | 55.61 ± 8.51<sup>b</sup> | 22.03 ± 6.23<sup>c</sup> | 12.8 ± 2.21<sup>c</sup> |
| Fertility (%) | 78.85 ± 4.82<sup>a</sup> | 24.76 ± 5.10<sup>b</sup> | 30.89 ± 10.67<sup>b</sup> | 19.32 ± 8.53<sup>b</sup> |
| Hatchability on FES (%) | 75.28 ± 5.17 | 60.00 ± 12.10 | 81.25 ± 13.15 | 90.00 ± 10.00 |
| No. of eggs incubated | 105 | 108 | 83 | 124 |

Values given are Mean±SE.
Figures bearing different superscripts in a row differ significantly ( $P<0.05$ ).
FES- Fertile egg set

## 4 DISCUSSION

In the present study a series of experiments were conducted to evaluate the effect of different cryoprotectants and diluents during Ghagus chicken semen cryopreservation. In this process semen cryopreservation protocols producing acceptable fertility were developed for this breed of chicken.

The DMSO and amide cryoprotectant DMA has been successfully used in cryopreservation of chicken semen protocols producing good fertility (Woelders et al., 2006; Santiago-Moreno et al., 2012; Herrera et al., 2005; Zhandi et al., 2017). Earlier, high fertility using 4% DMSO was obtained in another line maintained in the Institute farm (Pranay Kumar et al., 2019). In the present study the DMSO level was reduced to 2% based on preliminary in vitro studies, however, no fertility was obtained. In the present study DMA did not produce any fertile egg and is similar to an earlier report where DMA was used at 9% in Nicobari chicken (Shanmugam et al., 2018).

Ethylene glycol has been used for cryopreserving chicken semen and thawed at 5°C or 37°C (Mphaphathi et al., 2016; Miranda et al., 2018; Olexikova et al., 2019). The studies have reported only in vitro results and fertility trails were not conducted. Woelders et al. (2006) have reported use of EG as cryoprotectant during chicken semen cryopreservation where it was found to yield lower post-thaw motility and viability, however, the thawing conditions are not known. Ethylene glycol has also been used for storing chicken semen as pellets or in glass ampoules and there was no clear advantage over other cryoprotectants in terms of fertilizing ability (Wishart, 2001). The present study reports both in vitro and fertility results of experiments using EG as cryoprotectant. The interesting finding of this study was the influence of the thawing temperature on fertility outcome. Ethylene glycol cryopreserved semen when thawed at 5°C did not result in any fertility whereas semen thawed at 37°C resulted in 18-48% fertility. The fertility results were not affected by the semen diluent. Miranda et al. (2017) have hypothesized based on the in vitro results that due to the low molecular weight, greater membrane permeability and affinity towards sperm plasma membrane EG act as a better cryoprotectant when thawed at lower temperature. However, in the present study it was observed that semen cryopreserved with EG when thawed at 37°C rather than 5°C and inseminated produced higher fertility. It is not possible to compare results between these studies since fertility trails were not reported by Miranda et al. (2017). The results of the Experiments 2 and 3 indicate that thawing of cryopreserved semen is a critical point in the semen cryopreservation protocol. The in vitro results in the two experiments were almost similar or the values are lower in Experiment 3 where thawing was done at 37°C, however, better fertility was obtained. The thawing temperature has induced changes at cellular or molecular levels that are worth studying in further experiments.

Chicken semen cryopreserved in straws with DMF gave better post thaw motility parameters and fertility (Ehling et al., 2002). Higher post-thaw motility parameters were reported for semen cryopreserved with DMF and thawed at 5°C. Chicken semen cryopreserved only with DMF in plastic vials or straws produced higher fertility (Chalah et al. 1999; Chuaychu-noo et al., 2017; Thananurak et al., 2017, 2020). A higher live sperm, mitochondrial activity and lower acrosome damage was obtained in Korean native chicken semen cryopreserved with DMF (Choi et al., 2013). However, semen vitrified using DMF as a cryoprotectant resulted in negligible fertility (Shanmugam and Mahapatra, 2019). In the present report using DMF as cryoprotectant resulted in fertility ranging from 19 to 30%. Based on the fertility reports from these studies it may be concluded that DMF helps in cryopreserving sperm when packed in straws.

The composition of the diluent has been reported to influence the post-thaw in vitro semen parameters (Nabi et al., 2016; Rakha et al., 2016; Roushdy et al., 2014). The effectiveness of a particular diluent in these reports was based on in vitro parameters and no information is available about the fertility. Laboratory assays are only suggestive and cannot determine the fertilizing capacity of a semen sample, therefore, to know the fertility outcome from cryopreserved semen adequate number of inseminations is required (Graham and Mocé, 2005). The fertility after insemination with DMF cryopreserved semen was lower in a PVP based diluent compared to other diluents (Thananurak et al., 2017). In our study no difference in fertility was observed between PVP present or absent diluents in semen cryopreserved using DMF. Even in other experiments in our study it was observed that differences existed in post-thaw in vitro parameters between the semen diluents but not in fertility.

Acrosome reaction is an important parameter to be considered during semen cryopreservation. A low acrosome intact post-thaw sample will not produce fertile eggs, however, having higher post-thaw acrosome intact sperm will not guarantee fertility either. This could be observed in the first two experiments of this study. Induction of acrosome reaction in cryopreserved semen has been shown to be affected by type of birds/flock (Lemoine et al., 2011). Nevertheless, evaluation of acrosome integrity in post-thaw sperm may be used as a screening test along with other simple in vitro parameters such as motility to eliminate the cryoprotectant/protocol before proceeding further in the process of semen cryopreservation for any particular breed/line.

In conclusion, results from this study indicate that cryoprotectants and not diluents affect the fertility obtained from cryopreserved semen and the thawing temperature has been found to influence the success of cryopreservation protocol.

## AUTHOR CONTRIBUTIONS

SM designed the experiment, collected the samples, analysed the data and wrote the manuscript. RM designed the experiment and participated in the writing of the manuscript.

## DATA AVAILABILITY

Data for this study can be obtained from the corresponding author upon request.

## REFERENCES

Blesbois, E. (2011). Freezing avian semen. Avian Biology Research, 4, 44–50.

Burrows, W.H., & Quinn, J.P. (1937). The collection of spermatozoa from the domestic fowl and turkey. Poultry Science, 16, 19–24.

Campbell, R.G., Hancock, J.L., & Rothschild, L. (1953). Counting live and dead bull spermatozoa. Journal of Experimental Biology, 30, 44–49.

Chalah, T., Seigneurin, F., Blesbois, E., & Brillard, J.P. (1999). In vitro comparison of fowl sperm viability in ejaculates frozen by three different techniques and relationship with subsequent fertility in vivo. Cryobiology, 39, 185–191.

Choi, J.S., Shin, D-B., Ko, Y-G., Do, Y-J., Byun, M., Park, S-B., Seong, H-H., Kim, H., Kong, I-K., & Kim, SW. (2013). Effects of kinds of cryoprotectants on the characteristics of frozen fowl semen. Korean Journal of Poultry Science, 40, 171–178.

Chuaychu-noo, N., Thananurak, P., Chankitisakul, V., & Vongpralu, T. (2017). Supplementing rooster sperm with Cholesterol-Loaded-Cyclodextrin improves fertility after cryopreservation. Cryobiology, 74, 8–12.

Ehling, C., Taylor, U., Baulain, U., Weigend, S., Henning, M., & Rath, D. (2012). Cryopreservation of semen from genetic resource chicken lines. Agriculture and Forestry Research, 62, 151–58.

Graham, J.K., & Mocé, E. (2005). Fertility evaluation of frozen/thawed semen. Theriogenology, 64, 492–504.

Haunshi, S., Shanmugam, M., Rajkumar, U., Padhi, M.K., & Niranjan, M. (2015). Characterization of Ghagus breed vis-a-vis PD–4 birds for production, adaptability, semen and egg quality traits. Indian Journal of Animal Science, 85, 1338–1342.

Herrera, J.A., Quintana, J.A., López, M.A., Betancourt, M., & Fierro, R. (2005). Individual cryopreservation with dimethyl sulfoxide and polyvinylpyrrolidone of ejaculates and pooled semen of three avian species. Archives of Andrology, 51, 353–360.

Lake, P.E., Buckland, R.B., & Ravie, O. (1980). Effect of glycerol on the viability of fowl spermatozoa-implications for its use when freezing semen. Cryo-Letters, 1, 299–304.

Lake, P.E., & Ravie, O. (1984). An exploration of cryoprotective compounds for fowl spermatozoa. British Poultry Science, 25, 145–150.

Lemoine, M., Mignon-Grasteaue, S., Grasseau, I., Magistrini, M., & Blesbois, E. (2011). Ability of chicken spermatozoa to undergo acrosome reaction after liquid storage or cryopreservation. Theriogenology, 75, 122–130.

Long, J.A. (2006). Avian sperm cryopreservation: what are the biological challenges? Poultry Science, 85, 232–236.

Miranda, M., Kulíková, B., Vašíček, J., Olexiková, L., Iaffaldano, N., & Chrenek, P. (2018). Effect of cryoprotectants and thawing temperatures on chicken sperm quality. Reproduction in Domestic Animals, 53, 93–100.

Mphaphathi, M.L., Seshoka, M.M., Luseba, D., Sutherland, B., & Nedambale, T.L. (2016). The characterisation and cryopreservation of Venda chicken semen. Asian Pacific Journal of Reproduction, 5, 132–139.

Nabi, M.M., Kohram, H., Zhandi, M., Mehrabani-Yeganeh, H., Sharideh, H., Zare-Shahaneh, A., & Esmaili, V. (2016). Comparative evaluation of Nabi and Beltsville extenders for cryopreservation of rooster semen. Cryobiology, 72, 47–52.

Olexikova, L., Miranda, M., Kulikova, B., Baláži, A., & Chrenek, P. (2019). Cryodamage of plasma membrane and acrosome region in chicken sperm. Anatomia Histologia Embryologia, 48, 33–39.

Pope, C.E., Zhang, Y.Z., & Dresser, B.L. (1991). A simple staining method for evaluating acrosomal status of cat spermatozoa. Journal of Zoo and Wildlife Medicine, 22, 87–95.

Pranay Kumar, K., Swathi, B., & Shanmugam, M. (2019). Effect of supplementing L-Glycine and L-Carnitine on post thaw semen parameters and fertility in chicken. Slovak Journal of Animal Science, 52, 1–8.

Rakha, B.A., Ansari, M.S., Akhter, S., Hussain, I., & Blesbois, E. (2016). Cryopreservation of Indian red jungle fowl (*Gallus gallus murghi*) semen. Animal Reproduction Science, 174, 45–55.

Roushdy, Kh., El-Sherbieny, M.A., Abd El-Gany, F.A., & EL-Sayed, M.A. (2014). Semen cryopreservation for two local chicken strains as a tool for conservation of Egyptian local genetic resources. Egyptian Poultry Science Journal, 34, 607–618.

Santiago-Moreno, J., Castaño, C., Toledano-Díaz, A., Coloma, M.A., López-Sebastián, A., Prieto, M.T., & Campo, J. L. (2012). Cryoprotective and contraceptive properties of egg yolk as an additive in rooster sperm diluents. Cryobiology, 65, 230–234.

Sasaki, K., Tatsumi, T., Tsutsui, M., Niinomi, T., Imai, T., Naito, M., Tajima, A., & Nishi, Y., (2010). A method for cryopreserving semen from Yakido roosters using n-methylacetamide as a cryoprotective agent. The Journal of Poultry Science, 47, 297–301.

Shanmugam, M., & Mahapatra, R.K. (2019). Pellet method of semen cryopreservation: Effect of cryoprotectants, semen diluents and chicken lines. Brazilian Archives of Biology and Technology, 62, e19180188. DOI: 10.1590/1678-4324-2019180188.

Shanmugam, M., Pranay Kumar, K., Mahapatra, R.K., & Anand Laxmi, N. (2018). Effect of different cryoprotectants on post-thaw semen parameters and fertility in Nicobari chicken. Indian Journal of Poultry Science, 53, 208–211.

Silversides, F.G., Purdy, P.H., & Blackburn, H.D. (2012). Comparative costs of programmes to conserve chicken genetic variation based on maintaining living populations or storing cryopreserved material. British Poultry Science, 53, 599–607.

Thananurak, P., Chuaychu-noo, N., Thelie, A., Phasuk, Y., Vongpralub, T., & Blesbois, E. (2020). Different concentrations of cysteamine, ergothioneine, and serine modulate quality and fertilizing ability of cryopreserved chicken sperm. Poultry Science, 99, 1185–1198.

Thananurak, P., Chuaychu-noo, N., & Vongpralub, T. (2017). Freezability and fertility of Thai native chicken semen in different diluents. Thai Journal of Veterinary Medicine, 47, 551–556.

Wishart, G.J. (2001). The cryopreservation of germplasm in domestic and non-domestic birds. In Watson PF, Holt WV. Editors. Cryobanking the Genetic Resource: Wildlife Conservation for the Future? London: Taylor and Francis, p 183–185.

Woelders, H., Zuidberg, C.A., & Hiemstra, S.J. (2006). Animal genetic resources conservation in the Netherlands and Europe: poultry perspective. Poultry Science, 85, 216–222.

Zhandi, M., Ansari, M., Roknabadi, P., Zare Shahneh, A., & Sharafi, M. (2017). Orally administered Chrysin improves post-thawed sperm quality and fertility of rooster. Reproduction in Domestic Animals, 52, 1004–1010.

